# A single-cell transcriptomic comparison between small intestinal neuroendocrine tumors and their progenitor

**DOI:** 10.1101/2025.02.10.637596

**Authors:** Fredrik Axling, Elham Barazeghi, Per Hellman, Olov Norlén, Samuel Backman, Peter Stålberg

## Abstract

Several studies have attempted to find the initiating drivers of small intestinal neuroendocrine tumors (SI-NET) development and the molecular mechanisms driving progression and metastatic spread. For gene expression studies using bulk microarrays and RNA sequencing, researchers commonly use normal intestinal mucosa as a control. The intestine is made up of several different cell types, and using bulk RNA-seq may generate findings that reflect factors other than the tumor transformation. This could potentially contribute to the lack of discoverable treatments and prevention strategies for SI-NETs. We performed scRNA-seq on tissue from two patients that had tumor resection surgery and separated the EC cells from the normal intestinal mucosa and used specific markers to compare them against SI-NET tumors cells. We pinpointed new genes associated with chr18 haploinsufficiency but also others with loss or gain of expression that have not previously been associated with SI-NETs, and which could be potential targets for further functional genomic studies.

## Introduction

Small intestinal neuroendocrine tumors (SI-NETs), which are believed to originate from enterochromaffin (EC) cells in the gastrointestinal tract (de Herder 2005; Sei et al. 2016; Yao et al. 2008), are often slow-growing and small in size, with an annual incidence of 1 per 100,000. Most patients are diagnosed as stage IV with distant metastases (Modlin et al. 2014). Results from studies on epigenetic and genetic alterations in SI-NETs are scarce, and extensive DNA analysis using short-read sequencing and array-based methods has revealed a stable tumor genome, with frequent loss of chromosome 18 but no recurrent small mutations (Banck et al. 2013), with the exception of *CDKN1B*, which is only found mutated in 8% of cases (Crona et al. 2015; Francis et al. 2013). Several studies (Clift et al. 2019) have attempted to find the pathological origin of SI-NET upregulation and the underlying molecular mechanisms important for their progression and metastatic spread. For this purpose, normal intestinal mucosa is commonly used as a control for gene expression studies using bulk microarrays and RNA sequencing. Bulk analyses have become a mainstay of molecular biology research but are limited by the heterogeneity of tissues, which consist of multiple cell types, whose average gene expression is measured in bulk sequencing experiments (Li and Wang 2021). Single-cell RNA sequencing (scRNA-seq) has emerged as a powerful tool to study the gene expression patterns of individual constituent cells and has been successfully used to clarify the cellular origins of different forms of adult and pediatric kidney cancer (Young et al. 2018). Such an approach is highly useful for SI-NETs, which are thought to arise from a rare cell population in the intestinal epithelium, as the intestinal epithelium is made up of several different cell types (Elmentaite et al. 2021; Hickey et al. 2023), not only the SI-NET progenitor EC cell. Using bulk RNA-sequencing will generate results that are not specific to the SI-NET phenotype but rather reflect, e.g., the differences between enterochromaffin cells and healthy enterocytes. The paucity of relevant control samples may be one of the causes of the lack of translatable discoveries for targeted treatment strategies for SI-NETs.

To overcome this limitation, we have applied scRNA-seq to isolate the EC cells in the normal intestinal mucosa using specific cell markers and compare these against surgically resected SI-NET tumors. Initial efforts were focused on the development of protocols for obtaining viable suspensions of single cells from surgical specimens and required well-timed coordination between the operating team, the pathology lab, the research lab, and the core facility. In the present study, we aimed to compare the transcriptomes of SI-NET cells to the normal EC cell in order to identify driver genes and gene modules with aberrant expression specific to the SI-NET malignancy.

## Materials and methods

### Ethics statement

Informed written consent was obtained from all patients for tissue use and disease study, and ethical approval was secured from the Regional Ethical Review Board in Uppsala (EPN 2011/375). This research was carried out in compliance with the Declaration of Helsinki.

### Sample preparation, library construction and scRNA sequencing

During the gross pathological examination of surgical specimens from two patients undergoing surgical resection of SI-NETs, a representative piece of tumor tissue and a portion of adjacent normal bowel mucosa were taken for analysis. The tissue was minced, washed with Ham’s F-10 medium, and centrifuged to separate cells from the medium. Cells were transferred to a collagenase solution, followed by incubation at 37 °C for 2 hours, and filtered through a sterile stainless steel or nylon mesh to separate the dispersed cells and tissue fragments from the larger pieces (Birch 1989). Cells were then centrifuged and washed with EGTA solution, followed by Percoll treatment (Smedsrød and Pertoft 1985) to obtain a single-cell suspension. Sequencing libraries were prepared using the Chromium Single Cell 3’ Reagent Kit v3 (Zheng et al. 2017) (10X Genomics) in accordance with the manufacturer’s instructions. The resulting libraries were sequenced on one NovaSeq 6000 SP flow cell. The Cell Ranger Single-Cell Suite 7.2.0 was used to demultiplex and identify the assigned barcodes for each of the primary tumors and healthy adjacent tissue.

### External datasets for validation

A validation scRNA-seq dataset was *in silico* created using data from two different published studies: “Healthy small intestine from single-cell transcriptome analysis reveals differential nutrient absorption functions in the human intestine” (Wang et al. 2020) and primary SI-NET sample from “Comparative single-cell RNA sequencing (scRNA-seq) reveals liver metastasis-specific targets in a patient with small intestinal neuroendocrine cancer” (Rao et al. 2020). For bulk gene expression validation, two gene array expression datasets consisting of primary SI-NET tumor and normal mucosa tissue (Andersson et al. 2016; Leja et al. 2009) were merged using ComBat (Johnson, Li, and Rabinovic 2007) for batch correction. Followed by a logistic regression model applied to the remaining batches with a forward stepwise selection (FSS) algorithm to classify the batch-corrected samples. A Kolmogorov-Smirnov test was used for assessing distribution, and a Wilcoxon rank-sum test was used for differential expression. Overlapping enteroendocrine specificity was performed using the human protein atlas single-cell type transcriptomics map dataset (Karlsson et al. 2021).

### Bioinformatics processing and data analysis

Cell Ranger 7.2.0 (10x Genomics) was used with transcriptome reference 2020-A Human GRCh38 (GENCODE v32/Ensembl 98) to process raw sequencing samples from both the internal and external scRNA-seq datasets to generate count matrices, which were used as input for Seurat v5 (Butler et al. 2018) for downstream processing and analysis. The internal samples were quality controlled by discarding cells that contained less than 200 or more than 9000 features and had >90% mitochondrial gene expression. The external primary SI-NET sample was filtered by cells containing less than 250 or more than 5000 features with >10% mitochondrial gene content, and cells in the external small intestine samples were discarded if they contained less than 250 or more than 5000 features with >30% mitochondrial genes. EC and SI-NET cells were then isolated in both the internal and external samples using unsupervised clustering and specific cell markers, derived from the combination of the following markers: *CHGB, CHGA*, and *ADGRG4*. Other cell types were identified using the following markers: Fibroblast: *COL1A1, PDGFRA, and ACTA2*. Enterocyte: *APOA1, FABP6, ALPI, SI*, and *KRT20*. Leukocyte: *CD4, CD8A, CD3G*, and *KLRD1*. B cell: *CD79A* and *MS4A1*. Mast cell: *CPA3*. Monocyte: *CD14*. Foveolar cell: *IL33*. Goblet cell: *FCGBP, SPINK4, REP15*, and *MUC2*. Tuft cell: *AVIL, POU2F3, GFI1B*, and *TRPM5*. Stem cell: *LGR5, RGMB, SMOC2*, and *ASCL2*. Paneth cell: *LYZ*. Transit-amplifying cell: *PCNA, TOP2A*, and *MCM5*. NormalizeData and ScaleData, followed by IntegrateLayers, were used to merge cells derived from EC and SI-NET origins in each healthy and tumor sample into an internal discovery and an external validation dataset. Differential gene expression (DGE) between EC and SI-NET cells was then investigated in the discovery and validation datasets using FindMarkers with the Mann–Whitney U-test with Bonferroni correction (p-value) for multiple testing, and genes not overlapping between the two datasets were discarded. The STRING Database (Szklarczyk et al. 2023) with Cytoscape (Shannon et al. 2003) was used for producing interaction networks between differentially expressed genes. Functional gene enrichment analysis was performed using g:Profiler (Reimand et al. 2007).

## Results

### Cell type identification and specific gene expression differences

We performed single-cell RNA sequencing on tumor tissue and adjacent normal intestinal mucosa derived from two patients extracted during surgical resection of SI-NETs using the 10X Genomics Single Cell 3’ v3 chromium platform, yielding single-cell transcriptomes of 13086 cells. Unsupervised clustering and cell marker profiles were used for characterizing cells in the tumor and normal mucosa samples, followed by statistical comparison.

By comparing EC cells with SI-NET cells, our analysis revealed 741 significant (p ≤ 0.05) differentially expressed genes. To validate our novel scRNA-seq dataset, we *in silico* generated an external validation dataset. This was done using two datasets consisting of scRNA-sequenced normal intestine and a SI-NET tumor, applying the same approach as previously described. EC and SI-NET cells were isolated in the same manner from the external validation dataset and statistically compared with the resulting 1921 (p ≤ 0.05) significantly differentially expressed genes. Significant genes between the discovery and validation datasets were then compared, and genes without overlap were excluded from further analyses, resulting in a final comparison of 457 significant genes. Out of the 457 significant genes, 115 genes were upregulated and 342 were downregulated in tumor samples. Cells in the discovery and validation datasets can be distinguished in the Uniform Manifold Approximation and Projection (UMAP) plots in figure 1.

**Figure 1:**
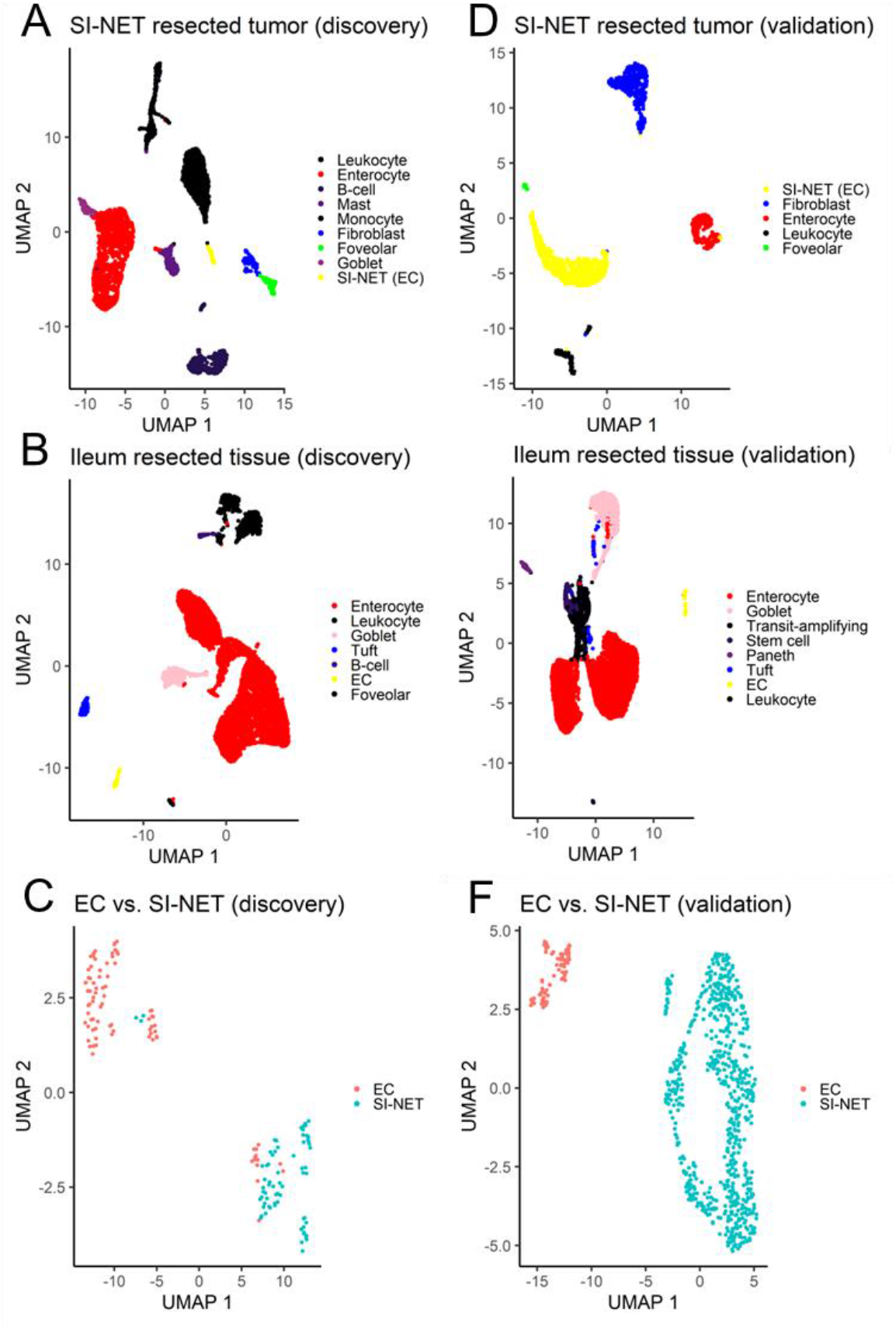
UMAP plots represent cells from the internal discovery dataset and the external validation dataset. A: Discovery SI-NET resected tumor cells (n = 4249), B: Discovery Ileum resected tissue cells (n = 6688), C: Discovery Enterochromaffin (EC) (n = 84) and SI-NET isolated cells (n = 69). D: Validation SI-NET resected tumor cells (n = 1373), E: Validation Ileum resected tissue cells (n = 7843), and F: Validation Enterochromaffin (EC) (n = 87) and SI-NET isolated cells (n = 660). Cell types: Enterocyte, Goblet, Fibroblast, Transit-amplifying, Stem, Tuft, Paneth, Enterochromaffin (EC), Small Intestinal Neuroendocrine Tumor (SI-NET), Foveolar, Monocyte, Mast, and B-cell.

### Gene expression methodology comparison

We compared overlap for our significant genes against earlier gene array expression data consisting of primary SI-NET and normal intestinal mucosa (Andersson et al. 2016; Leja et al. 2009). Re-analyzing the gene array datasets revealed 6685 significant genes between primary SI-NET and normal mucosa, but only 216 genes (67 tumor-upregulated and 149 tumor-downregulated genes) preserved the same trajectory of expression between the scRNA-seq and bulk gene array dataset.

### Gene interaction network

We proceeded to cluster the significant genes that passed validation using a gene expression network approach with a minimum required interaction score of 0.9 and isolated 103 interconnected neighboring genes. In tumor cells, 16 of the 103 connected genes showed upregulation, while 87 displayed downregulation (Figure 2).

**Figure 2:**
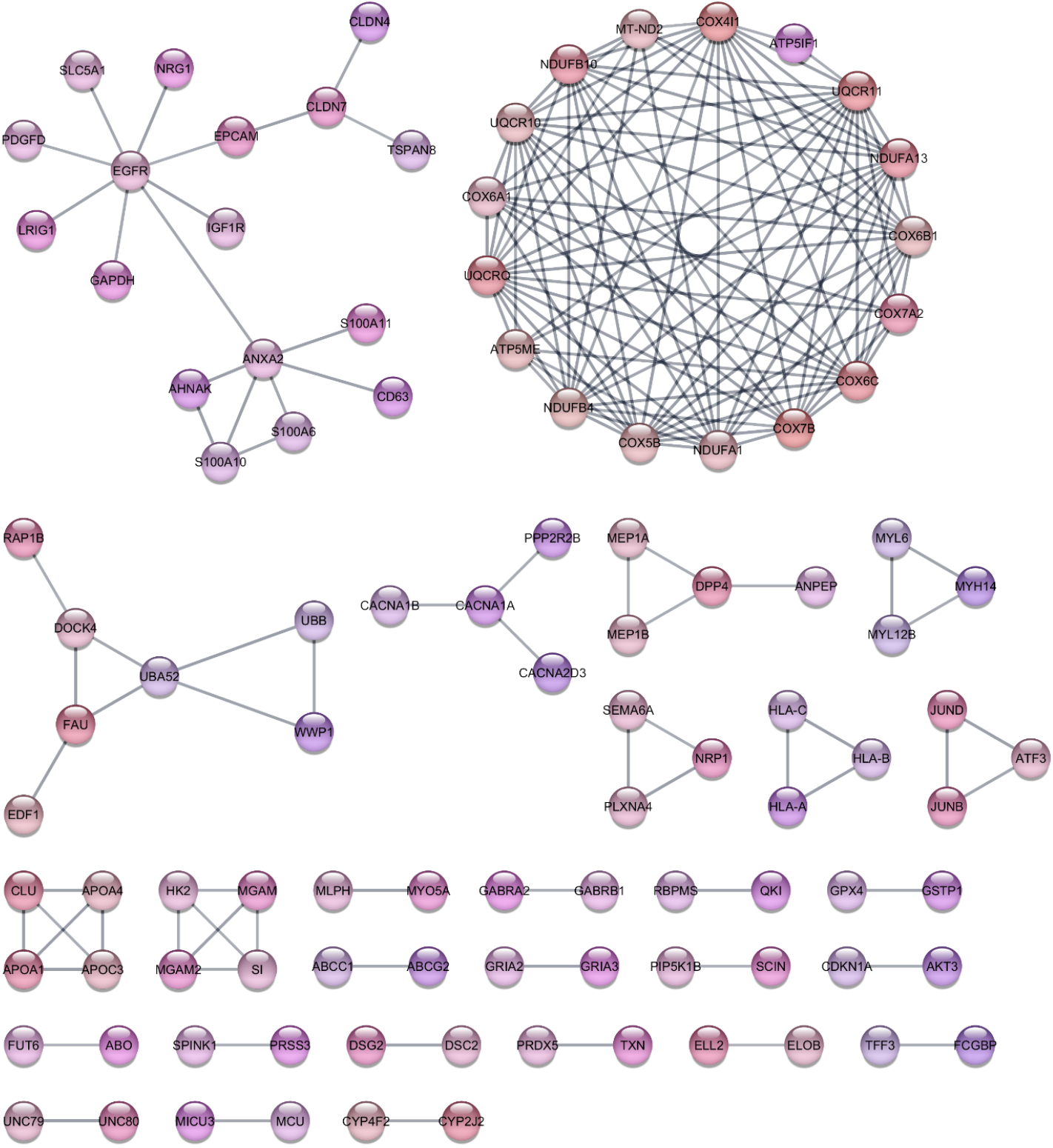
The gene expression network inputs from 457 significant genes with a minimum required interaction score of 0.9 and at least one connected neighboring gene, resulting in 103 genes with neighboring interactions.

We conducted a gene enrichment analysis of the genes involved in the network, and the top driver terms revealed for the SI-NET upregulated genes were AMPA glutamate receptor activity, amyloid-beta binding, monoatomic ion channel complex, AMPA glutamate receptor complex, and neuron projection, and the SI-NET downregulated terms can be observed in figure 3. We linked the Reactome pathway results to TP53 signaling and tyrosine kinase receptor (ERRB4) signaling.

**Figure 3:**
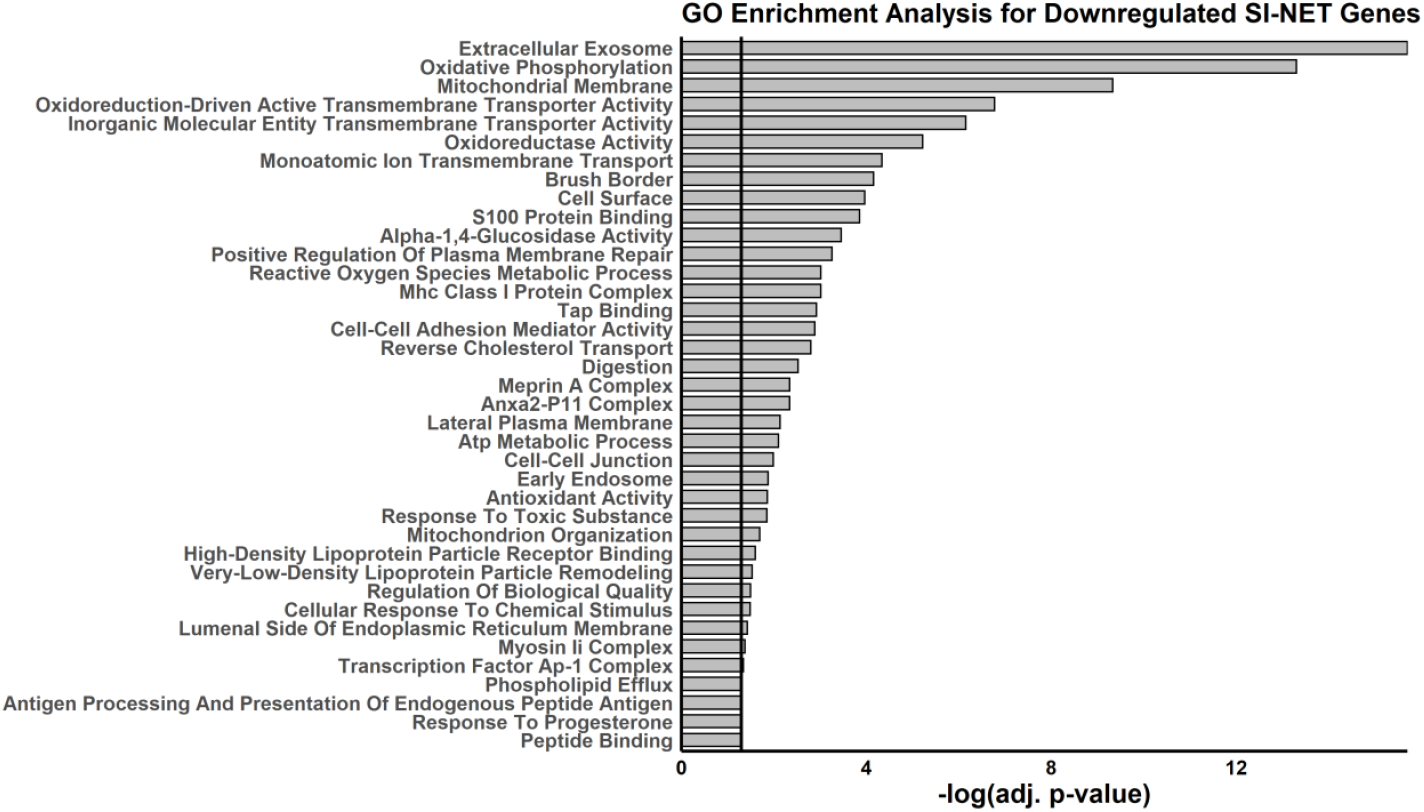
Gene ontology (GO) analysis of the 87 SI-NET downregulated genes. The black line indicates p-value threshold (adj. p-value = 0.05).

### Loss of chromosome 18 heterozygosity

As loss of heterozygosity on chromosome 18 is a common mutational event in SI-NETs, we also examined genes located on chromosome 18. We found 13 genes with a significant downregulation of expression in tumor cells (Table 1).

**Table 1:**
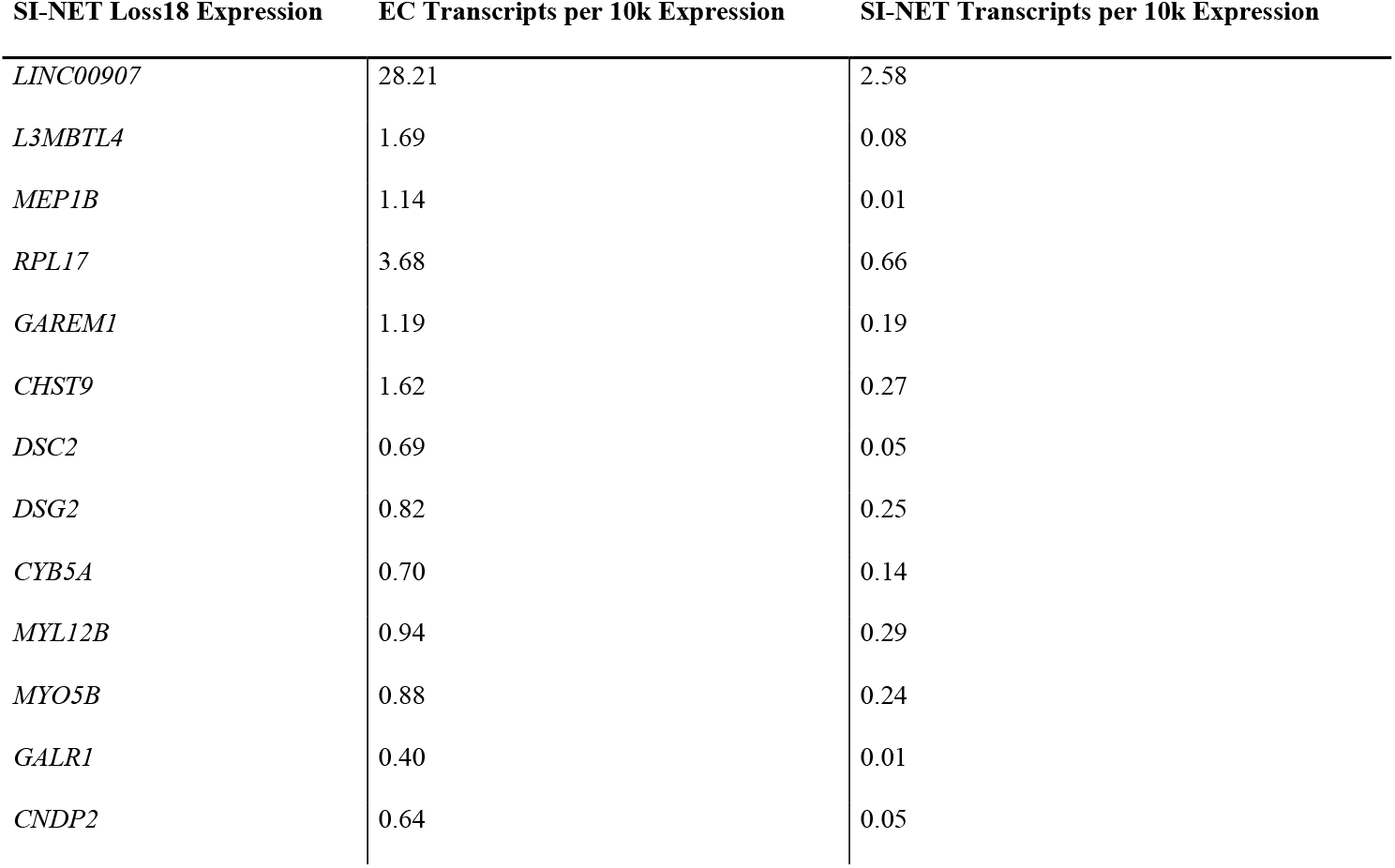
Genes located on chromosome 18 with downregulation of expression in SI-NET cells.

### SI-NET tumorigenesis impact on gene expression differentiation

We proceeded to investigate genes with major downregulation in tumor cells, and in total, we could detect 20 genes that had abolished expression (expressed in <1.5% of cells). However, only *ISL1, ONECUT3, PPP1R1C, LCN15, NTS*, and *PLXNA4* had specificity for enteroendocrine expression in the small intestine when compared against single-cell expression data in the human protein atlas single-cell type transcriptomics map. The expression of these six downregulated genes can be observed in figure 4.

**Figure 4:**
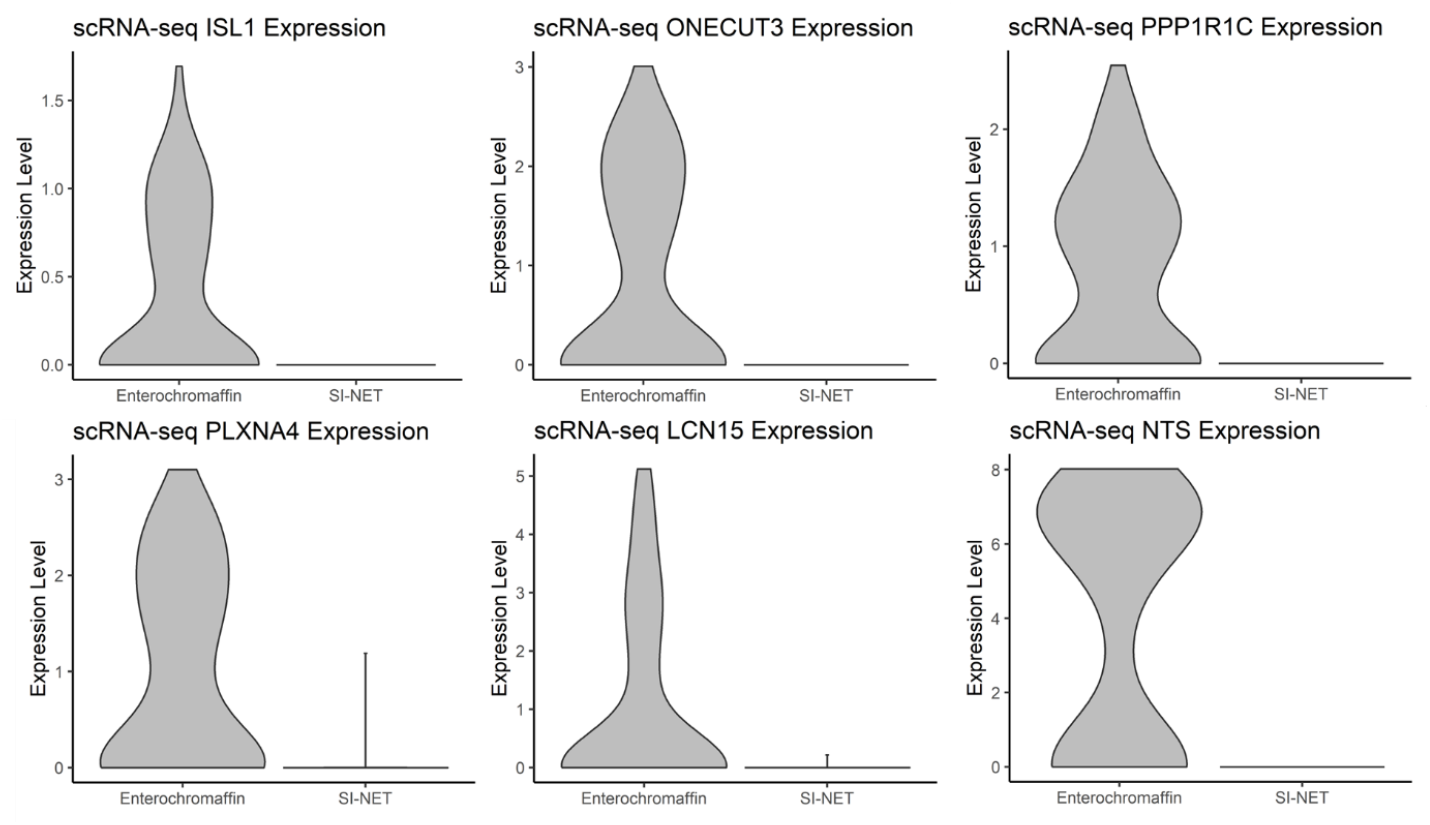
Expression patterns of the six genes with verified expression in enteroendocrine cells were revealed to be downregulated in the SI-NET tumor cells.

In contrast, 32 genes with no expression in EC cells but with presence in SI-NET cells could be confirmed in the human protein atlas database to lack enteroendocrine expression. *ADAMTS9-AS2, NPSR1-AS1, LINC02388*, and *LINC02484* could not be verified due to their lack of presence in the human protein atlas database. Further experiments will be needed to investigate the possible role of these dysregulated genes (Table 2).

**Table 2:** Human protein atlas-verified genes (n = 38) with gain or loss of expression in SI-NET cells. LoE = loss of expression; GoE = gain of expression.

| SI-NET LoE | SI-NET GoE | SI-NET GoE |
| --- | --- | --- |
| *ISL1* | *FBXL7* | *ARHGAP6* |
| *LCN15* | *MYT1L* | *ANKS1B* |
| *NTS* | *UBE2E2* | *SLC6A15* |
| *ONECUT3* | *ZMAT4* | *DENND2A* |
| *PLXNA4* | *GRIK3* | *UNC5D* |
| *PPP1R1C* | *NMNAT2* | *ADAM19* |
|  | *HS3ST4* | *ATP8A2* |
|  | *HPN* | *SYT14* |
|  | *ABLIM3* | *ATCAY* |
|  | *UBE2QL1* | *ZBTB46* |
|  | *ZNF626* | *DACH2* |
|  | *ELAVL3* | *GLRA3* |
|  | *MICU3* | *FGF13* |
|  | *ADAM12* | *QKI* |
|  | *TCEAL2* | *MMP16* |
|  | *XKR4* | *GRIA2* |

## Discussion

Until now, EC-related studies have mainly been focused on their role and function in the gastrointestinal tract (Anon 2017; Beumer et al. 2020; Singh et al. 2024). While their comparison against SI-NETs has been conducted using bulk-tissue methodologies (Samsom et al. 2019), others have focused on the comparison between primary and metastatic lesions (Rao et al. 2020) and intertumoral differences (Hoffman et al. 2023) using single-cell technologies. Here we present the first single-cell sequencing study comparison between the small intestinal neuroendocrine tumor cell and its progenitor enterochromaffin cell. Until 10x Genomics released their 10X Chromium FLEX platform, applications of single-cell techniques required fresh viable cells, and this proved to be a challenge to obtain from surgically resected SI-NET specimens, which is evident in the comparatively low cell number and relatively high degree of mitochondrial gene expression. This had a significant impact on the number and quality of the tumor cells included in the analysis; in order to mitigate this and to decrease the risk of false-positive findings, we performed a two-step validation where we first compared our results for overlapping expression against an external validation dataset. This resulted in a final list of 457 overlapping significant genes, and we proceeded to pinpoint enriched differences using gene interaction networks and gene ontology analysis for expression differences between EC and SI-NET. This showed numerous enriched differences between the two cell types, revealing functions associated with tumor relevance in the receptor tyrosine kinase and TP53 pathways, as well as effects corresponding to cellular respiratory mechanisms. We also observed numerous cell morphology associations, including cell junctions, cell adhesion, plasma membrane, and cytoskeleton. However, this enrichment is not unexpected in a tumor versus normal cell comparison, but an interesting observation is that the majority of the genes associated with these pathways are downregulated in the tumor cells.

SI-NETs experience recurrent loss of chromosome 18 heterozygosity affecting the transcription capabilities of its tumor suppressor gene *SMAD4* (Banck et al. 2013), and thus inducing SMAD4-haploinsufficiency (Hofving et al. 2021). We could distinguish 13 genes located on chromosome 18 with downregulated expression in the tumor cells. The long, non-coding RNA (lncRNA) LINC00907 was the most downregulated, but also the adenylyl cyclase inhibitor *GALR1* (Galaninreceptor 1) and GRB2-associated and regulator of MAPK protein 1, *GAREM1*, are noteworthy. LncRNAs possess tissue-specific expression and exhibit phenotypic regulated expression (Guttman et al. 2009) with a number of lncRNAs transcriptionally controlled by important oncogenes or tumor suppressors (Huarte et al. 2010; Zheng et al. 2014). LINC00907’s function is not clear, but it is associated with adenosquamous prostate carcinoma and chromosome 18q deletion syndrome (Rappaport et al. 2017).

We summarized a subset of 38 genes that passed overlapping validation with either abolished or initiated expression in SI-NET cells and had been second-step validated for the presence of enteroendocrine expression in the human protein atlas database. We compared the expression of these 38 genes in our GOT1 and CNDT2.5 xenografts (unpublished data), as these cell lines can be valuable for conducting functional genomic studies in SI-NETs. The following genes had no or very low expression in both cell line xenografts: *NTS, LCN15, ONECUT3, ZMAT4, HS3ST4, HPN, UBE2QL1, TCEAL2, ADAM19, MICU3, NMNAT2*, and *DACH2*.

Out of the 38 genes, six genes with loss of expression in tumor cells; a subset of these was shown to be of particular interest, with *ISL1* (encoding for LIM Homeobox 1, an insulin gene enhancer region binder protein, among other targets) already being reported in earlier studies as lacking expression by immunohistochemistry analysis in SI-NETs. (Tran, Sherman, and Howe 2020; Xiang, Malik, and Zhang 2023). On the other hand, little is known about *LCN15*, which encodes lipocalin 15. Our results showed that *NTS* exhibited the strongest expression differences, with 15 log2 fold changes, a difference that is also evident in our bulk gene expression comparison. *NTS* expression comparison in our CNDT2.5 and GOT1 xenografts revealed no or very little expression of NTS, and these results demonstrate that NTS expressions are downregulated in SI-NETs.

Thanks to non-cancerous tissue investigations, NTS has been shown to display a vital role in the promotion of wound healing (Mouritzen et al. 2018). Its receptors have been linked to the Wnt/β-catenin signaling pathway in pancreatic NETs, glioblastoma, and hepatocellular carcinoma, and upregulation leads to tumor growth (Kim, Liu, et al. 2015; Xiao et al. 2017; Ye et al. 2016). Most likely, the same biological mechanisms upregulating wound healing also promote tumor growth (Liu et al. 2022; Whyte, Smith, and Helms 2012). NTS and its high-affinity receptor NTSR1 are most extensively characterized in the context of gastrointestinal cancers. Both are overexpressed in gastric and colon cancer, and this overexpression correlates well with poor prognosis (Christou et al. 2020). Four different non-intestinal NET cell lines have been shown to express neurotensin (*NTS*) on both the transcript and protein levels (Kim, Li, et al. 2015), and when upregulated, stimulate cell proliferation and growth (Kim, Liu, et al. 2015). However, more research is needed to find out what role NTS might play in SI-NETs and how its role in tumor growth and cell proliferation differs from those seen in colorectal and stomach cancer.

We used a very low cut-off (expressed in less than 1.5% of cells) to decide which genes are either losing or gaining expression in SI-NETs. This means that there are probably more genes that should be looked into among the differentially expressed genes. Moreover, relative changes in gene expression, in addition to the upregulation of silenced genes and the complete silencing of expressed genes, may play a crucial role in tumor development. Therefore, we urge further screening for associations of tumor relevance in our significant genes to possibly cull more candidates for functional studies. This study provided us with a glimpse of potential genes responsible for or being affected during the tumorigenesis and growth of small intestinal neuroendocrine tumors and which would be of interest to investigate by functional genomic studies in SI-NET cells.

In conclusion, by using single-cell RNA sequencing, we were able to distinguish unexplored differences between SI-NET and its progenitor cell, and we identified a few important genes for future studies.

## Data

We obtained gene array expression data for bulk expression comparison from GEO accession no. GSE9576 and GSM1591697. Single-cell transcriptome data for the external validation dataset was obtained from GSE125970 for the ileum and GSE140312 for the primary small intestine neuroendocrine tumor.

## Acknowledgments

This study was funded by grants from the Swedish Cancer Foundation and governmental funding of clinical research within the Swedish National Health Service (ALF). The authors are grateful to the Endocrine Surgical Unit at the Uppsala University Hospital for assistance. Single-cell sequencing was performed by the SNP&SEQ Technology Platform in Uppsala. The facility is part of the National Genomics Infrastructure (NGI) Sweden and Science for Life Laboratory. The SNP&SEQ Platform is also supported by the Swedish Research Council and the Knut and Alice Wallenberg Foundation.

